# Play vocalizations induce affective bias in reward learning and memory in orangutans

**DOI:** 10.64898/2026.05.22.726256

**Authors:** Daan W. Laméris, Colin Allen, Heidi Lyn, Christopher F. Martin, Emma K. Nelson, Ximena J. Nelson, Alex Taylor, Erica A. Cartmill

## Abstract

Affective biases, or shifts in learning and decision-making driven by affective states, are central to human cognition. Translational studies in rodents have shown that pharmacological and stress manipulations alter reward valuation, but how positive social signals shape these processes remains poorly understood. Here, we adapted the Affective Bias Test (ABT), a translational rodent assay of affective biases in depression and antidepressant therapy, to ask whether positive social signals influence reward valuation in orangutans (*Pongo* spp.). We first validated the task by manipulating reward magnitude during learning and found that participants showed a significant preference for substrates previously associated with higher reward value, confirming sensitivity to reward valuation. We then tested whether play vocalizations influence reward learning by presenting either conspecific play vocalizations or control sounds prior to learning reward-substrate associations, with identical reward values. In subsequent preference tests, orangutans showed a significant choice bias for substrates previously associated with play vocalization recordings. These findings demonstrate that orangutans experience positive affective states when hearing conspecific play vocalizations, as indicated by affective biases in reward valuation and memory.

## Introduction

Affective states play a central role in shaping how organisms perceive, respond to, learn from, and remember experiences. Emotionally salient events are more readily encoded and retrieved than neutral ones [1], a phenomenon known as an affective bias. In humans, positive mood enhances learning, increases attribution of value to experiences, and produces lasting biases in associative memory [2]. As such, beyond their hedonic quality, positive affective states are associated with adaptive cognitive mechanisms that reflect how environmental experiences are valued, and shape subsequent decision-making in ways that promote exploration, reward seeking, and continued interaction with positive contexts [3–6]. While the importance of affective biases in human decision-making is broadly acknowledged, the evolutionary origins of affective biases in reward valuation and associative memory is largely unknown [7].

Affective states arise not only from direct experience of the world; emotional expressions of conspecifics can induce corresponding affective states in others through emotional contagion [8]. Laughter and its accompanying facial expressions occupy a central role in human social interactions as sources of powerful emotional contagion [9], orienting cognitive resources toward positive social stimuli [10] and promoting motivation to approach others, social affiliation, and the development of cognitive and socio-emotional skills [3]. These patterns are not unique to humans, but are shared with great apes, where play faces readily attract attention [11], and are rapidly mimicked during playful interactions [12–14]. Play-associated vocalizations homologous to human laughter show similar contagious effects [15] and have been linked to enhanced optimism in bonobos [16]. However, it is unknown whether such vocalizations influence the encoding and valuation of reward-associated memories, a fundamental aspect of associative learning. This leaves a critical gap in our understanding of the role of vocalizations during play: social play is not simply nor exclusively enjoyable, but represents a critical context for development of cognitive, motor, and social competence across animals [17,18]. If play-associated vocalizations induce positive states and bias how co-occurring experiences are encoded and valued, they may represent a functional mechanism through which positive affect facilitates learning and the development of competence.

To address this question, we adapted and applied the Affective Bias Test (ABT) to ask whether hearing conspecific play vocalizations would impact reward valuation in orangutans (*Pongo* spp.). The ABT was originally developed as a translational assay of affective state-induced biases in reward learning and memory in rodents [19]. The test predicts that if an affective state during learning shifts cognitive valuation of a reward, this should lead to preference biases in a subsequent choice task, even when objective reward values are equal across conditions. The ABT has direct translational parallels with human affective neuroscience [20] and has been extensively validated using pharmacological and environmental manipulations in rodents: antidepressant and pro-depressant drug treatments induce respective positive and negative affective biases [19,21,22]. Similarly, exogenous hormone treatments influence affective bias, with oestradiol and the oxytocin analogue carbetocin inducing positive biases, while progesterone and testosterone (in males) induce negative biases [23]. Blocking androgen furthermore resulted in a positive bias effect and blocking oestrogen produced a negative bias [23]. Manipulations of environmental variables can likewise influence affective biases, with positive biases reported in rodents after enriched housing conditions [24], social play sessions [21], and tickling protocols [25], while restraint stress causes a negative bias [21].

Here, we tested whether communicative vocal signals produced in positive social contexts could induce positive affective biases in great apes using a modified ABT task. We first validated the task in orangutans by manipulating reward magnitude during learning, confirming sensitivity to reward valuation. We then asked whether hearing conspecific play vocalizations induced positive biases in the encoding and valuation of rewards. If play vocalizations induce positive affect that impacts cognitive valuation, orangutans should show a preference bias for substrates previously associated with play vocalizations, despite equal objective reward values across conditions. This would establish a functional mechanism by which positive communicative signals add value to experiences.

## Methods

### Ethics statement

All sessions were conducted using positive reinforcement and conform to the guidelines of the Ex-situ Program (EEP), formulated by the European Association of Zoos and Aquaria (EAZA), and complied with the ASAB guidelines [26]. Participation in test sessions was voluntary and were conducted in moments where individuals were separated for husbandry training. This study was approved by the Ethics Committee of the Royal Zoological Society of Antwerp, Indianapolis Zoo, and Indiana University (24-033-1) and conformed to the American Society of Primatologists Principles for the Ethical Treatment of Non-Human Primates.

### Subject and housing conditions

Six orangutans (two females and four males; mean age = 31.7 years; range = 8-51 years old) housed in two zoological institutions (Indianapolis Zoo, USA, n = 1; and Zoo Planckendael, Belgium, n = 5) participated in the current study (Table S1). The Indianapolis Zoo housed twelve orangutans at the Simon Skjodt International Orangutan Center, which comprises interconnected indoor and outdoor enclosures, including an indoor dayroom with 50ft high ceilings and multiple outdoor yards connected by an elevated trail system reaching up to 80ft that promotes species-typical arboreal locomotion. Planckendael Zoo housed five orangutans in a tropical greenhouse consisting of three main enclosures, each connected to outdoor islands. Both institutions followed flexible managed fission-fusion dynamics based on husbandry demands, aimed at mimicking the natural social structure of orangutans. All orangutans had multiple feeding moments per day consisting of a variety of vegetables, supplemented with browse and nutritionally balanced biscuits and had *ad libitum* access to water.

### Materials and Task Design

We adapted and applied the ABT [19,27] for orangutans. The task used a two-alternative forced choice design and was conducted in one of the home enclosures of the orangutans, out of view of visitors. The setup consisted of two plywood platforms, each equipped with a bowl-like structure formed by raised ridges on three sides, leaving the fourth side open to allow easy manual access by the apes (Figure 1A). On each trial, one bowl was baited with a preferred food reward (e.g., almond, cashew nut) fully concealed beneath a substrate (e.g., shredded paper), while the other bowl contained a different substrate (e.g., yarn pieces) but no reward. The reward was not visible and required active manual searching to be located. For each trial, both platforms were presented simultaneously, positioned parallel to the front of the enclosure, allowing manual access to the bowls. A choice was defined as the first instance of searching behavior, i.e., the displacement of substrate material with a finger. Once a participant began exploring one bowl, the other platform was immediately withdrawn so that only one substrate could be explored per trial. Location of the rewarded substrate was randomized across trials, and the identity of the target substrate was counterbalanced between participants. Novel substrate combinations were used for each phase (training, Reward Learning Assay, Affective Bias Test, Table S2).

**Figure 1:**
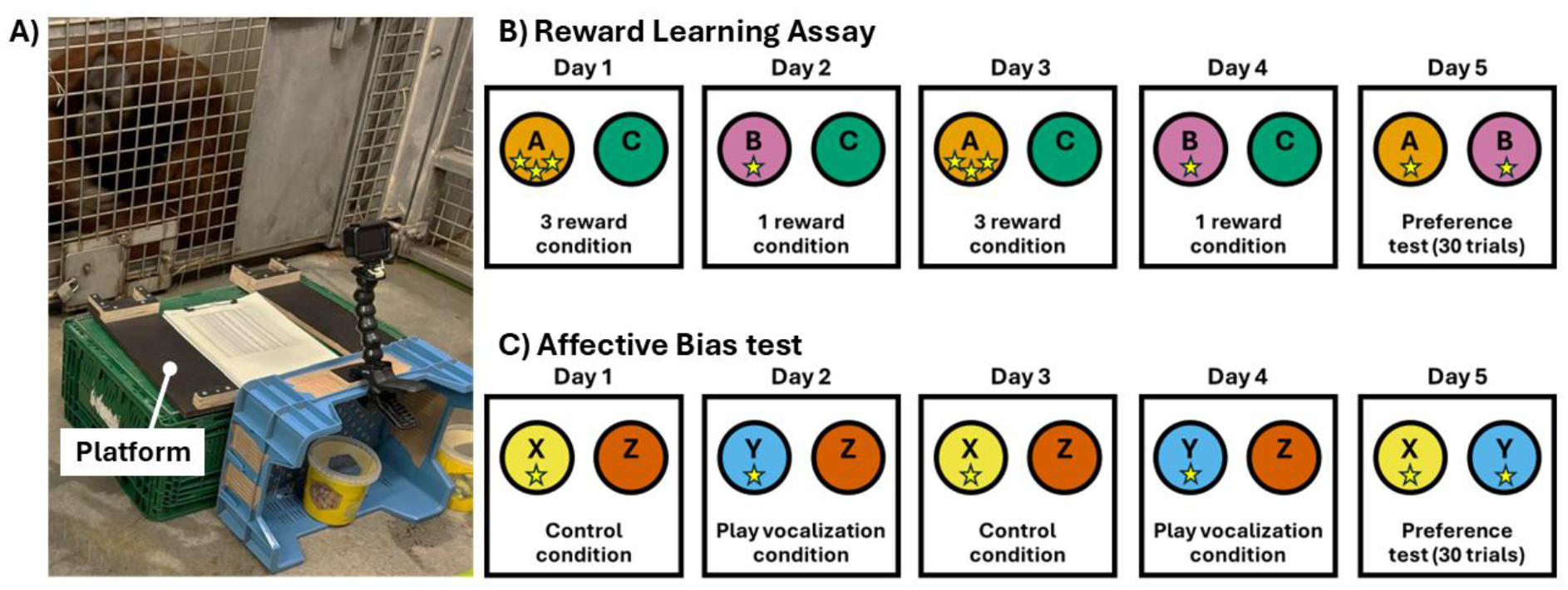
Experimental setup in Zoo Planckendael (A) and conceptual overview of the Reward Learning Assay (B) and Affective Bias Test (C). In both experiments, orangutans learned associations between distinct substrates and a food reward (indicated by the star symbol) across four pairing sessions (Days 1-4), followed by a preference test (Day 5). In the Reward Learning Assay, one substrate (e.g., substrate A) was paired with three rewards and the other (e.g., substrate B) with one reward. In the Affective Bias Test, reward quantity was held constant between the substrates (e.g., substrates X and Y) but learning occurred either following conspecific play vocalization playback, or a control sound. The order of conditions was counterbalanced across participants.

### Training

Orangutans were first familiarized with a simplified version of the task in which rewards were readily visible (i.e., bowl without substrates) for ten trials. We then introduced substrate coverage over ten trials until the reward was fully concealed.

Once familiarized, we presented the orangutans with a discrimination session of 30 trials in which the two bowls were filled with two distinct substrates and a single reward was buried in one of them behind a barrier, so the participant could not see the hiding location. Once the participant achieved six consecutive correct trials (choosing the substrate associated with the reward), the session ended. If they did not reach this criterion, the session was repeated. However, all subjects reached criterion on the first session. Randomization and counterbalancing procedures for substrates were as described above.

### General testing protocol

The general testing protocol for the Reward Learning Assay and Affective Bias Test involved associative learning between two different substrate-reward pairings, each learned across two sessions on alternating days (days 1 and 3 for one pairing, days 2 and 4 for the other, Figures 1B-C). During each session, the two platforms were presented with two different substrates: one substrate that was reward-paired, and the other as an unrewarded control. Each session consisted of a maximum of 30 trials, ending early if the participant achieved six consecutive correct choices. Control substrates were constant across the four pairing sessions and between apes. The two substrate-reward pairings were learned under control and experimental conditions. Any resulting preference bias between these pairings could therefore be attributed solely to the experimental manipulation induced at the time of learning. Affective biases during the experimental conditions were then quantified in a choice bias test the day after the last pairing session. During this choice bias test, we presented the orangutans with 30 trials of the two previously learned reward-paired substrates. To avoid new associative information, each substrate was baited with a single reward on a random one-third of trials, with the remaining trials unrewarded.

### Reward Learning Assay

To validate task sensitivity to reward valuation we conducted a Reward Learning Assay (RLA), after the training steps and prior to completing the Affective Bias Test. Instead of using an experimental condition to manipulate an affective bias, animals learned to associate one of the two reward-paired substrates with either a high or low value reward to test the effects of absolute reward value (Figure 1B).

Prior to the RLA, we completed a quantity preference test with the Planckendael orangutans. We presented the animals with three and one reward at the same time, allowing them to choose only one of the two options. All five Planckendael orangutans selected the larger quantity significantly above chance (p = 0.031, two-tailed binomial, Table S3).

During the RLA, one reward-paired substrate was associated with three reward items, and the other substrate was paired with one reward. On the fifth day, we completed the choice test, measuring the preference for either of the two substrates.

### Affective Bias Test

After the RLA, we conducted the Affective Bias Test (ABT) and tested whether hearing play vocalizations induced an affective bias. As such, prior to the two substrate-reward pairings, the orangutans listened to five min of conspecific play vocalizations, predominantly laughter and play squeaks [28], or a control sound. Recordings from unfamiliar subadult orangutans were compiled into a 2.5-min clip, repeated once to create a continuous 5-min stimulus. As a control sound, we used a volume matched, synthetic environmental wind noise [16]. Audio was played from two Logitech Z150 speakers connected to a laptop, with a consistent volume level across all sessions. The experimenter asked the orangutan basic medical husbandry commands (e.g., presenting body parts) throughout the playback and periodically gave a small fruit reward, approximately every 10-15 s, to motivate them to remain near the speakers. Immediately after the playback, the orangutan started pairing sessions as per the protocol, using new substrates and equal rewards between testing conditions (Figure 1C).

### Statistical Analysis

For the RLA and ABT, we calculated individual-level preference scores by expressing each participant’s proportion of choice towards the target substrate as a percentage and centering at zero (choice bias score = proportion x 100 - 50). Here, a score of zero reflects chance performance, positive scores reflect preference for the treatment-paired substrate, and negative scores a bias for the control-paired substrate. Individual-level significance was assessed using two-tailed binomial tests against an expected proportion of 0.5.

At the group level, we tested whether mean preference scores differed significantly from zero using one-sample t-tests using the stats package [29]. Given the small sample size inherent to great ape research, we could not reliably verify the normality assumption underlying the parametric t-test. We therefore additionally ran a bootstrapped t-test with 10,000 resamples. Bootstrapped p-values were derived as the proportion of resampled t-statistics equal to or more extreme than the observed t-statistic in absolute value. Effect sizes are reported as Cohen’s d with 95% confidence intervals using the effectsize package [30].

Learning curves across sessions were analyzed using a mixed-effects Cox proportional hazards model using the coxme package [31], with trials to criterion as the time variable and a binary indicator of criterion attainment as the event. Condition, session repetition, and presentation order were included as fixed effects alongside their two-way interactions. Participant was included as a random effect to account for repeated measures within individuals.

All analyses were done in RStudio version 2026.1.2.418 [32], and figures were created using the ggplot [33] and survminer packages [34].

## Results

### Reward Learning Assay

To validate task sensitivity to reward valuation, we tested whether orangutans showed a preference for substrates previously associated with a higher reward magnitude. Individual-level analyses revealed that four of six individuals showed a significant preference for the high-reward substrate (p < 0.05, Table S4). The remaining two orangutans did not show a preference for one of the two substrates (p > 0.5). At the group level, mean preference scores were significantly above zero (mean = 16.7 ± 13.0 SD, t(5) = 3.14, p = 0.026, bootstrapped p = 0.063, Cohen’s d = 1.28, 95% CI [0.14, 2.36], Figure 2), indicating a large overall effect of reward magnitude on substrate preference. We note that the bootstrapped p-value was marginally above the conventional threshold, reflecting the influence of small sample size and the single individual whose preference score was in the opposite direction.

**Figure 2:**
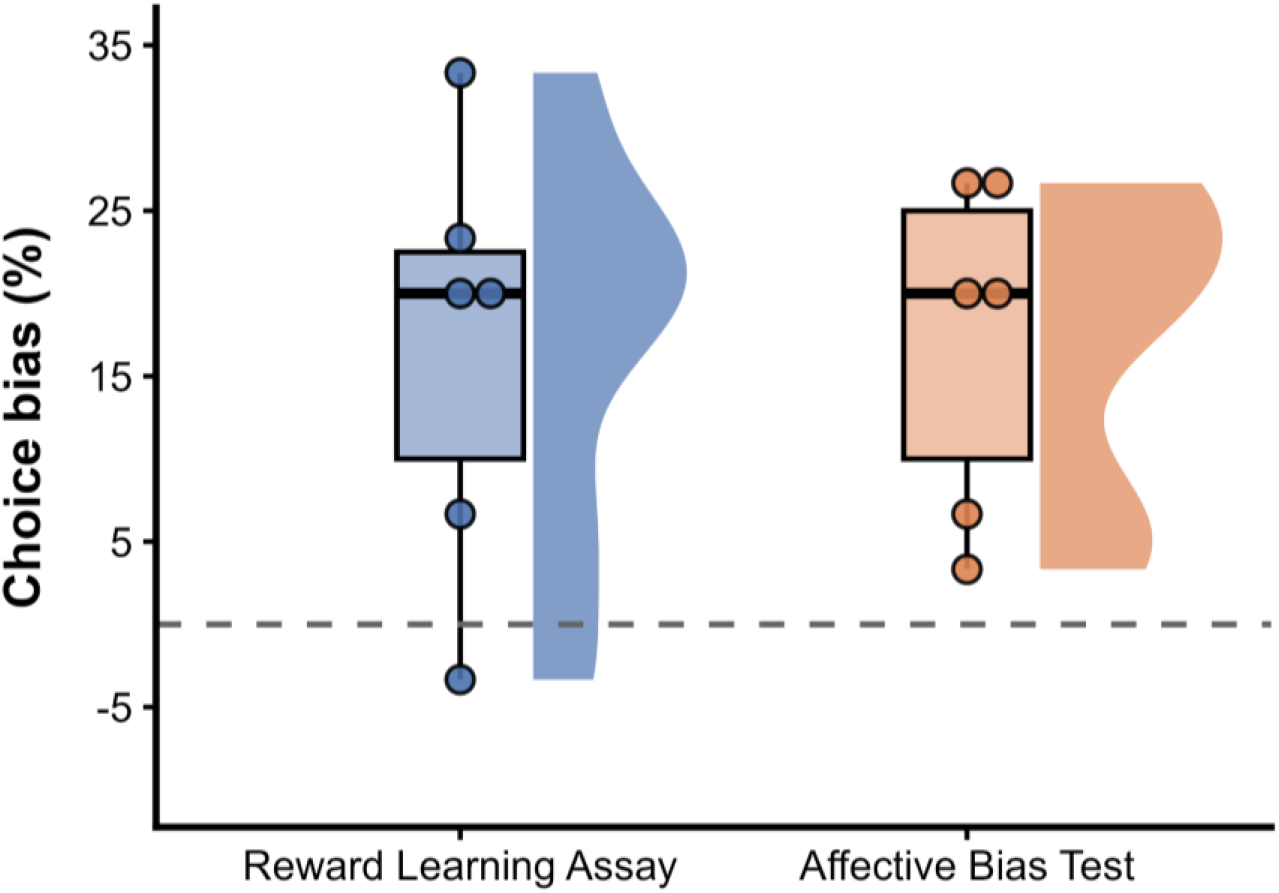
Choice bias scores for the Reward Learning Assay and Affective Bias Test. Choice bias is expressed as the percentage of choices toward the target substrate minus 50. Boxes represent the interquartile range with median, violins show the kernel density distribution, and dots represent the individual data points. Horizontal dashed line indicates chance performance.

To examine if reward magnitude influenced the rate of learning, we analyzed trials to criterion. There was no interaction effect between reward condition and session repetition (HR = 1.73, z = 0.54, p = 0.588) or between reward condition and presentation order (HR = 0.16, z = -1.77, p = 0.077) on learning speed. Reward condition alone also did not predict learning speed (HR = 1.90, z = 0.79, p = 0.429), indicating that the high and low reward substrates were learned at equivalent rates. Session repetition, however, significantly predicted faster criterion attainment (HR = 6.26, z = 2.43, p = 0.015), reflecting a familiarity effect whereby individuals reached criterion faster in repeat sessions. Presentation order did not predict learning speed (HR = 5.23, z = 1.61, p = 0.108).

### Affective Bias Test

We tested whether exposure to conspecific play vocalizations prior to learning induced an affective bias in reward valuation. Individual-level analyses revealed that all six individuals showed a preference score in the direction of the play vocalization-paired substrate, of whom four reached individual significance (p < 0.05, Table S5). At the group level, mean preference scores were significantly above zero (mean = 17.2 ± 10.0 SD, t(5) = 4.23, p = 0.008, bootstrapped p = 0.029, Cohen’s d = 1.73, 95% CI [0.39, 3.01], Figure 3), indicating a large effect of exposure on subsequent substrate preference.

**Figure 3:**
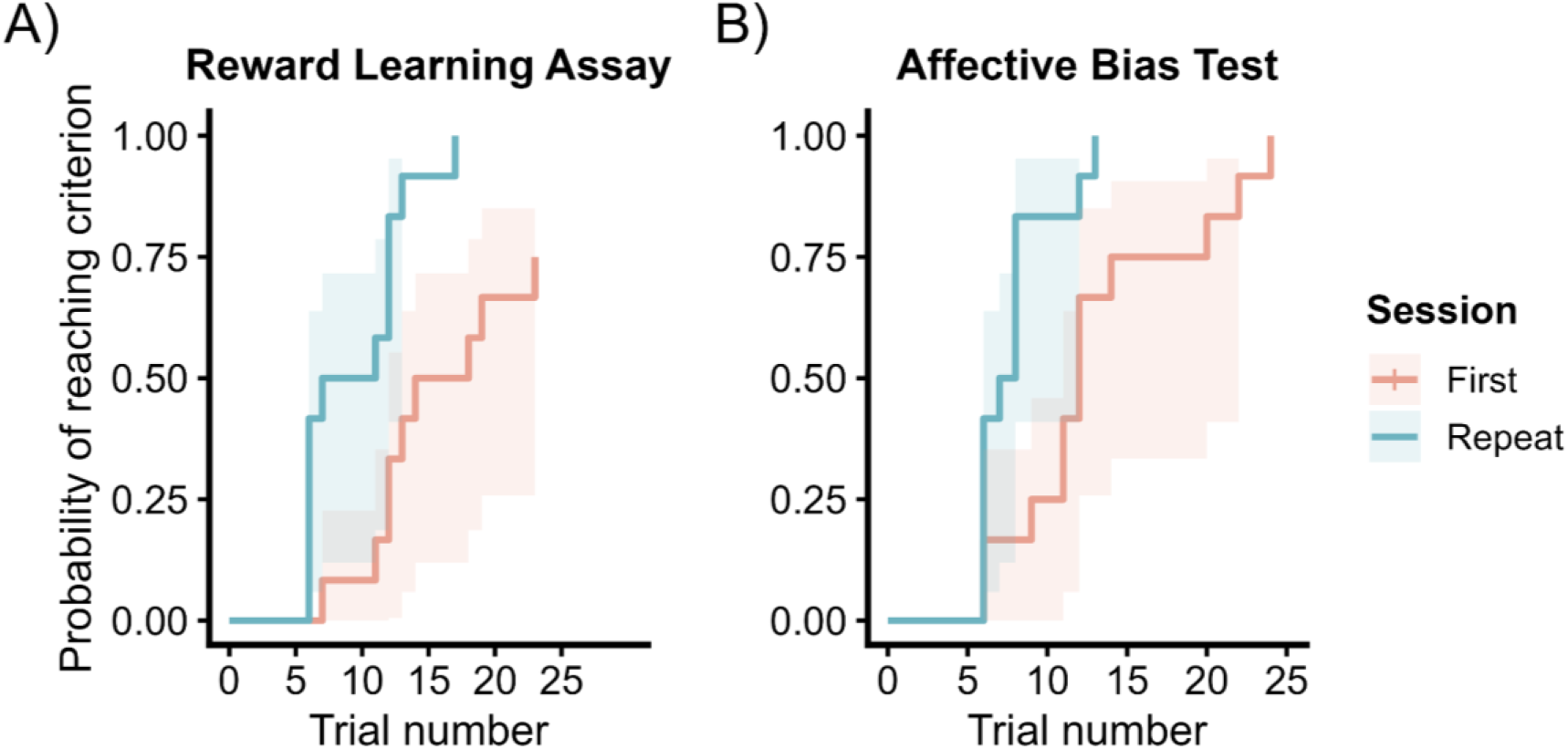
Cumulative probability of reaching learning criterion as a function of trial number for the first (orange) and repeat (blue) sessions, for the Reward Learning Assay (A) and Affective Bias Test (B). Shaded areas represent 95% confidence intervals.

With regard to learning rates, we did not find an interaction effect between condition and session repetition (HR = 2.81, z = 1.07, p = 0.283), or between condition and presentation order (HR = 1.86, z = 0.68, p = 0.495). Hearing play vocalizations or the control audio also did not predict learning speed (HR = 0.26, z = -1.43, p = 0.152), indicating that substrate-reward associations were acquired at similar rates. As in the RLA, session repetition significantly improved criterion attainment (HR = 4.15, z = 2.10, p = 0.036), confirming a consistent familiarity effect across both experiments. Presentation order similarly did not predict learning speed (HR = 2.80, z = 1.39, p = 0.165).

## Discussion

Our results suggest that hearing communicative signals of positive affect biases reward valuation and associative memory in orangutans. Importantly, neither actual reward value nor play vocalization impacted learning speed during training days, indicating that these effects impact the consolidation of reward value, rather than the speed of information acquisition. This is consistent with the hypothesis that positive affective states, here induced by conspecific play vocalizations, influence how experiences are encoded and subsequently valued. This raises the possibility that play vocalizations not only function as a social signal or as an expression of positive internal states, but potentially contribute to the role of play in learning and skill acquisition through enhanced encoding and valuation of co-occurring experiences.

The RLA provided important validation of the paradigm in orangutans by showing that the apes’ memories were sensitive to reward magnitude: when one substrate was associated with three rewards and the other with one, orangutans showed a significant preference for the substrate associated with the larger reward. Importantly, a similar preference emerged when reward quantity was held constant, but one substrate had been paired with conspecific play vocalization playback, indicating that audible positive vocalizations can create memory biases and influence later reward-based choices.

Our findings significantly broaden previous work showing that positive social or playful experiences can induce positive affective biases in animals. In rats, playful sessions produce positive affective shifts that are detectable in the ABT [21], and tickling by human handlers induces optimism in both an ambiguous cue task [35] and the ABT [25]. Importantly, previous manipulations involve direct playful experiences, whereas our study shows that hearing play-associated vocalizations is salient enough to bias cognitive responses. This adds to previous findings that hearing conspecific laughter increases optimism in a judgement bias task with rats [36] and bonobos [16]. Beyond expressing positive internal states, play-associated vocalizations may have divergent effects on cognitive functioning in receivers. That is, where judgement bias paradigms measure interpretation of ambiguous cues, the ABT measures how affective states bias the encoding of reward value during associative learning. The play vocalization-induced affective biases we observed reflect an effect on memory encoding and reward valuation, suggesting that play-associated vocalizations may have deeper consequences for cognitive functioning in receivers than previously acknowledged.

The observed affective bias is furthermore consistent with emotional contagion, the induction of a corresponding affective state in the receiver through exposure to another’s emotional expression [8,37]. Notably, we did not observe overt play vocalizations in response to the playbacks, suggesting that the effect was not mediated by behavioral contagion, providing experimental dissociation between emotional and behavioral contagion. While cases of behavioral contagion in orangutans, like scratch contagion [38], yawn contagion [39], and rapid facial mimicry during play [12], may reflect underlying emotional contagion, they are distinct phenomena. Our findings suggest that emotional contagion occurred in the absence of behavioral contagion and that the cognitive consequences of emotional contagion can be measured independently of any overt behavioral responses.

These findings fit within the broaden-and-build theory of positive affective states [3], which proposes that positive affect builds lasting cognitive resources. A growing body of comparative literature reports effects of positive signals on various cognitive processes ([10,11,40] but see [41]), suggesting that this is a conserved trait across great apes. Laughter is a phylogenetic homologous vocalization across the Hominidae [42], and the current findings provide novel insights in the functionality of this communicative signal with deep evolutionary roots in playful contexts. These findings are consistent with the view that cognitive mechanisms linking positive affect to memory encoding are shared across the Hominidae, mirroring well-documented effects of positive mood on learning and memory in humans [2,3].

In conclusion, our findings highlight that hearing conspecific play vocalizations biased orangutan reward valuation and memory in a way that is consistent with induced positive affective states. This strengthens the view that play vocalizations, beyond signalling positive affect in the expressor, may play a broader role in shaping cognition in receivers during social interactions.

## Acknowledgements

We are grateful to the staff of the Royal Zoological Society of Antwerp (RZSA) and the Indianapolis Zoo for their support in this study. Special thanks go to Marina Davila-Ross for generously providing the orangutan recordings, and the great ape keepers for their assistance throughout the study. Finally, we thank the orangutans who participated in this study.

## Author contributions

Conceptualization: DWL, EAC; Investigation: DWL, CFM, EKN; Methodology: DWL, EAC; Formal analysis: DWL, EAC; Writing – Original draft: DWL, EAC; Writing – Review & Editing: DWL, CA, HL, CFM, EKN, XJN, AT, EAC; Funding acquisition: CA, HL, ZJN, AT, EAC

## Data availability

Data and code will be made available at Zenodo upon publication.

## Supporting information

**Table S1:** Orangutan participant demographics.

| Participant | Sex | Date of birth | Species | Zoo |
| --- | --- | --- | --- | --- |
| Azy | Male | 14-12-1977 | Bornean x Sumatran hybrid | Indianapolis Zoo |
| Bagus | Male | 17-06-2002 | Sumatran | Zoo Planckendael |
| Ombak | Male | 04-03-2017 | Sumatran | Zoo Planckendael |
| Vilmos | Male | 09-12-2010 | Sumatran | Zoo Planckendael |
| Karo | Female | 08-12-1973 | Sumatran | Zoo Planckendael |
| Moni | Female | 07-12-1977 | Sumatran | Zoo Planckendael |

**Table S2:** Example combinations of substrates used.

| Substrate A | Substrate B | Substrate C | Location used |
| --- | --- | --- | --- |
| Paper towel squares | Leaf litter | Fabric pieces | Indianapolis Zoo |
| Construction paper (green-black mix) | Yarn (orange) | Crepe paper (neutral) | Indianapolis Zoo |
| Wood wool | Carton shreds | Hemp | Zoo Planckendael |
| Moss | Yarn (yellow-blue mix) | Earth | Zoo Planckendael |
| Paper confetti | Cotton | Saw dust | Zoo Planckendael |

**Table S3:** Result of the quantity preference testing.

| Participant | Score | P-value (one-tailed) |
| --- | --- | --- |
| Bagus | 10/12 | 0.019 |
| Moni | 17/20 | 0.001 |
| Vilmos | 9/10 | 0.011 |
| Ombak | 11/12 | 0.003 |
| Karo | 10/12 | 0.019 |

**Table S4:** Individual-level preference scores and corresponding two-tailed binomial test p-values for the Reward Learning Assay. Choice biases are calculated as (proportion x 100) - 50.

| Participant | Sum chosen<br>target substrate | Proportion chosen<br>target substrate | Choice<br>bias | Binomial test<br>p-value |
| --- | --- | --- | --- | --- |
| Azy | 14 | 0.467 | -3.333 | 0.855 |
| Bagus | 25 | 0.833 | 33.333 | 0.0003 |
| Karo | 21 | 0.700 | 20.000 | 0.043 |
| Moni | 21 | 0.700 | 20.000 | 0.043 |
| Ombak | 17 | 0.567 | 6.667 | 0.585 |
| Vilmos | 22 | 0.733 | 23.333 | 0.016 |

**Table S5:** Individual-level preference scores and corresponding two-tailed binomial test p-values for the Affective Bias Test. Choice biases are calculated as (proportion x 100) - 50.

| Participant | Sum chosen<br>target substrate | Proportion chosen<br>target substrate | Choice<br>bias | Binomial test<br>p-value |
| --- | --- | --- | --- | --- |
| Azy | 16 | 0.533 | 3.333 | 0.855 |
| Bagus | 17 | 0.567 | 6.667 | 0.585 |
| Karo | 21 | 0.700 | 20.000 | 0.043 |
| Moni | 21 | 0.700 | 20.000 | 0.043 |
| Ombak | 23 | 0.767 | 26.667 | 0.005 |
| Vilmos | 23 | 0.767 | 26.667 | 0.005 |

